# Embodied Latch Mechanism of the Mandible to Power at an Ultrahigh Speed in the Trap-jaw Ants *Odontomachus kuroiwae*

**DOI:** 10.1101/2022.12.05.519247

**Authors:** Hitoshi Aonuma, Keisuke Naniwa, Yasuhiro Sugimoto, Kyohsuke Ohkawara, Katsushi Kagaya

**Author notes:** Address for Correspondence: Hitoshi Aonuma, D.Sc., Graduate School of Science, Kobe University, 1-1 Rokkodai, Nada-ku, Kobe, Hyogo 657-8501, Japan. Department of Mechanical Engineering, Osaka University, Suita, Osaka 565-0871, Japan (Keisuke Naniwa).

## Abstract

Rapid movements of limbs and appendages, faster than those produced by simple muscle contraction alone, are generated through mechanical networks consisting of springs and latches. The latch plays a central role in these spring-loaded mechanisms, but the structural details of the latch are not always known. The mandibles of the trap-jaw ant *Odontomachus kuroiwae* closes the mandible extremely quickly to capture prey or to perform mandible-powered defensive jumps to avoid potential threats. The jump is mediated by a mechanical spring and latch system embodied in the mandible. An ant can strike the tip of the mandible onto the surface of an obstacle (prey, predator, or ground) in order to bounce its body away from potential threats. The angular velocity of the closing mandible was 2.3×10^4^ rad/s. Latching of the joint is a key mechanism to aid the storage of energy required to power the ballistic movements of the mandibles. We have identified the fine structure of two latch systems on the mandible forming a ‘ball-joint’ using an X-ray micro-computational tomography system (X-ray micro-CT) and X-ray live imaging with a synchrotron. Here we describe the surface of the inner section of the socket and a projection on the lip of the ball. The X-ray live imaging and movements of the 3D model show that the ball with a detent ridge slipped into a socket and over the socket ridge before snapping back at the groove edge. Our results give insight into the complex spring-latch systems that underpin ultra-fast movements in biological systems.

## Introduction

Speed is a crucial factor for animals to survive in their changing environments. During a prey-predator encounter, the predator and/or the prey moves quickly; the former to capture the prey and the latter to escape from the threat. The evolution of power-amplified rapid ballistic movements underpins survival for many animals in order that they can avoid potential threats and capture elusive prey. Many arthropods have evolved specialized appendages to produce quick movements (e.g. locust; (Heitler, 1974), dragonfly larva; (Pritchard, 1965), mantis shrimp; (Patek et al., 2004); snapping shrimp (Ritzmann, 1974; Versluis et al., 2000)) Longo et al. (2019) offered a principled foundation for the latch-mediated spring actuation of the animals. The speed of movement at velocities above the physiological limit of muscle contraction is produced and controlled by dynamically coupled, neuro-mechanical systems. The locust jump, for example, represents one of the best known quick movements used by animals to escape from the predatory threats (Burrows, 1995). The locust fully flexes the hindlegs to prepare for the jump and co-contracts the extensor and flexor tibia muscles. Then the extensor tibia muscles slowly contract to store energy in the extensor apodeme and exoskeleton. The jump is triggered by the inhibition of motor neurons and flexor muscles. An strage of energy storage before a quick movement of the labium is also found in the dragonfly larvae (Tanaka and Hisada, 1980). The extensor and flexor muscles of the labium are co-contracted before a quick extension. Relaxation of the flexor muscles and a hydraulic mechanism produce a predatory strike at a high speed (Tanaka and Hisada, 1980). Mantis shrimps store elastic energy in the exoskeleton to generate to power for a quick raptorial strike (Zack et al., 2009) where an exoskeletal spring mechanism with several components, such as a saddle or ventral bar, is thought to be a key (Patek et al., 2004).

Many ant genera employ so-called trap-jaw mechanisms that have independently evolved at least four times distantly related ant lineages and seven times within a single ant genus (Booher et al. 2021). Ants of the genus *Odontomachus* have long and powerful trap-jaws that function as a weapon when hunting (De la Mora et al., 2008). The ant closes the mandible extremely quickly to capture prey (Gronenberg et al., 1993; Gronenberg, 1996; Just and Gronenberg, 1999). Using the mandible, these trap-jaw ants also perform the jump so-called bouncer defense jumps (Patek et al., 2006; Spagna et al., 2009). More aggressive ants respond to unexpected tactile stimuli with defensive behavior and potentially show the mandible-powered jumps (Aonuma, 2020). The jump is thought to increase survival rate during predator-prey encounters (Larabee and Suarez, 2015). The trap-jaw ants approach prey by fully opening the mandible. The mandible has mechanoreceptive hairs that can detect prey and capture it at an ultra-fast speed. The sensory neurons originate from hairs and project to the suboesophageal ganglion to activate mandibular motor neurons (Gronenberg and Tautz, 1994; Gronenberg et al., 1993; Gronenberg, 1995). The key to the ultra-fast movements is thought to be a mechanism of the power amplification realized by the combination of a catapult mechanism and elastic energy storage in the apodeme and skeletal systems (Gronenberg, 1996). The ultra-fast movement of the mandible consists of 4 phases: (i) open, (ii) latch, (iii) load, and (iv) strike phase. The contraction of the mandible abductor muscles opens the mandible, and the fully opened mandible is latched to prepare for the loading phase. The contraction of the large adductor muscles initiates the loading phase, and the contraction of the trigger muscle releases the latch to promote a quick strike of the mandible (Gronenberg and Ehmer, 1996). The latching mechanism is crucial for the trap-jaw ants to power the mandible strike at an ultra-fast speed.

Here we performed X-ray micro-imaging to examine the structures of the mandibular joint that could function as latch components. X-ray micro-CT scanning and X-ray *in vivo* live imaging demonstrate that the mandibular joint of the trap-jaw ant genus *Odontomachus* is like a ball-joint with fine projections on the surface of the lip and the inner section of the socket.

## Materials and Methods

### Animals

Workers of the trap-jaw ants *Odontomachus kuroiwae* were mainly used in this study. Colonies of the trap-jaw ants *O. kuroiwae* were collected in Okinawa. The trap-jaw ants *Odontomachus monticola* (Emery 1892) collected in Osaka were also used to compare the performance of the mandible strike. The two species have been classified as different species (Yoshimura et al., 2007), however, since they are closely related their morphological characteristics are quite similar. The trap-jaw ants are mostly polygynous and their colonies each contain 3-4 queens, 100-300 workers and broods. Each colony was installed in an artificial plaster nest (200 mm×200 mm×40 mm) in a plastic case (600 mm×400 mm×200 mm) on a 14:10 hr. light and dark cycle (lights on at 6:00) at 25±2°C. They were fed a diet consisting of insect jelly (Marukan Co., Ltd, Osaka, Japan), cricket nymphs and water *ad libitum*. Notably, there are no distinctive differences between major and minor workers both in *O. kuroiwae* and *O. monticola*.

### Mandible powered defensive jumps

The trap-jaw ants of the genus *Odontomachus* show mandible powered jumps named bouncer defense (Patek et al., 2006; Spagna et al., 2009). We first analyzed the mandible powered jump in both *O. kuroiwae* and *O. monticola*. An ant was placed into an acrylic arena (450×450×200 mm). After a short rest (5min), the mandible-powered jump was introduced by the tactile stimulation using a steel rod (*φ*=8 mm). The ant responded to the approaching rod by turning toward it and opening its mandibles (Aonuma, 2020). Since the mandible has mechanosensory hairs that detect contact with an obstacle (Gronenberg et al., 1993), the tactile stimulation was applied to the sensory hairs. Mandible powered jumps were recorded using a pair of high-speed cameras that were placed at the top and one side of the arena (2,000 fps, HAS U2, DITECT, Tokyo, Japan). Each frame was stored in TIFF format for later analysis. The movement of the ant was tracked to measure jump velocity using an open-source video tracking software Kinovea (Ver 0.8.15, https://www.kinovea.org).

### Recording ultrafast movements of the mandibles

The movement of the mandibles was observed under a dissection microscope (SZX-12, Olympus, Tokyo, Japan). An ant was anesthetized using carbon dioxide gas and mounted on a plastic platform using wax (GC corporation, Tokyo, Japan). Ultrafast movement of the mandibles was elicited by directing an air puff to the head and recorded using a high-speed camera (10,000-60,000 fps, HAS D71, DITECT, Tokyo, Japan) attached to the dissection microscope and each frame was stored in TIFF format for later analysis. The movement of the tip of the mandible was tracked to measure the angular velocity using the video tracking software Kinovea.

### Kinematic analysis during the opening phase

Angular velocities of mandible movements during the opening phase were measured using high-speed imaging. The tips of both mandibles were filmed at 1,000 fps and tracked to measure the angular velocity using the video tracking software Kinovea.

To analyze the latch kinematics of the opening phase of the mandible movements, we constructed statistical models to estimate the parameters using the probabilistic programming language Stan through the R package “rstan”. We assumed that the pivot remained stationary during the opening phase, and then estimated the trajectories of the tip point of the mandible by fitting circles using the pivot point and the time series of the angular position. From the resulting time series of angular position, we calculated the instantaneous velocity using a five-point stencil approximation formula:

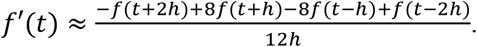

This formula calculates the derivative of *f* with respect to time, *f*’(t), at a given time point *t*, using the function values at two points on either side of *t*, separated by a distance of *h*. The formula provides an approximation to the true derivative value, and we used it to calculate the instantaneous velocity.

Our model for the angular velocity had the two components, (1) the overall value *μ* and (2) a specific value to each strike *μ_strike_*. The formula for AngularVelocity[*n*] was given by:

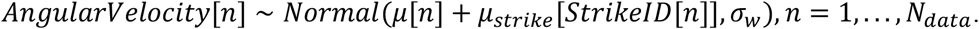

To constrain the value of *μ* at the current angular position to be similar to the previous position, we used a normal distribution with a parameter *σ_t_*, which acted as a soft constraint. Specifically, we used the formula:

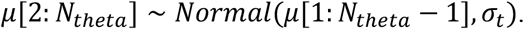

The strike specific value can be generated from this distribution with a mean value specific to each individual animal *μ_animal_* and the standard deviation *σ_1_*:

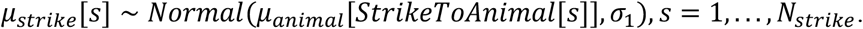

The *μ_animal_* assumed to be generated a normal distribution with the mean 0 and the parameter σ_0_:

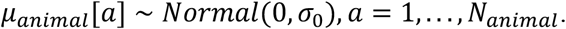

### X-ray micro-computed tomography imaging

The *O. kuroiwae* ants were anesthetized using carbon dioxide and were fixed with alcoholic Bouin’s fixative in a refrigerator (3°C) for 2 days. After fixation, samples were dehydrated with ethanol series (70% twice, 80%, 90% and 100% for 60 min each) and then stained using 1% iodine diluted in 100% ethanol for 2 days. Staining was used to enhance contrast of each ant tissue during X-ray micro-computed tomography (micro-CT) scanning. After staining, samples were transferred to liquid *t*-butyl alcohol at 40°C. Samples were then placed in a refrigerator (3°C) to solidify *t*-butyl alcohol quickly. They were then freeze-dried by using a handmade freeze dry system composed of a vacuum evaporator (PX-52, Yamato Ltd., Japan) with a cold alcohol trap (H2SO5, AS ONE, Japan). All chemicals were obtained from Kanto Chemical Co. (Tokyo, Japan).

Samples were scanned on an X-ray micro-CT system (inspeXio SMX-100CT, Shimadzu Corporation, Kyoto, Japan) whose X-ray source was operated at 40 kV and 100 μA. Images were reconstructed with a voxel size of 2-3 μm. Image reconstruction and rendering were carried out using the software VGStudio MAX (ver. 2.2.6 125, Volume Graphics, Heidelberg, Germany) and amira (ver. 2019.1, Thermo Scientific, Waltham, USA). Some of the CT images were converted into STL files to print out 3D models using a 3D printer (REPLICATOR2, Makerbot, New York, U.S.A.). The 3D models were magnified 110 times and used to examine the movement of the mandibular joint.

### X-ray *in vivo* live imaging

The X-ray *in vivo* live imaging was performed using a synchrotron at SPring-8 in Hyogo Japan. Live *O. kuroiwae* ants were scanned in Hutch 3 of beamline 20B2, located over 200m from the storage ring whose X-ray source was operated at 33 keV. The ant was mounted on a plastic rod platform (*ϕ* =5 mm) using wax (GC corporation, Tokyo, Japan) to avoid head movements during the experiment. It was placed between the X-ray source and the detector. Live images were recorded using a Hamamatsu phosphor charge-coupled device (CCD) detector (C4742-95HR) with an active area of 24 × 15.7 mm^2^ (100 fps). All images were recorded with a spatial resolution of approximately 2.74 μm. Beam inhomogeneities and detector artefacts were corrected with custom software using images of the direct beam and detector dark current. All images scanned were saved in TIFF format and reconstructed using image processing software Image J (NIH, ver. 2.0.0).

## Results

Initial analysis focused on the 3D structure of the mandibular join to gain a better understanding of the latching mechanism of the ant *O. kuroiwae*. The morphology of *O. kuroiwae* was similar to that of *O. monticola*. The body color of both of these species was both brown, however the *O. monticola* was much darker than *O. kuroiwae*. The body mass of the *O. kuroiwae* was smaller than that of *O. monticola*. The matured workers of *O. kuroiwae* were 11.2 ± 1.13 mg (mean ± SD, N=10) and that of *O. monticola* was 14.6 ± 1.62 mg (N=8) (Table 1). The single mandible mass of the *O. kuroiwae* was 261 ± 3.51 μg (mean ± SD, n=20) and *O. monticola* was 360 ± 4.37 μg (n=16).

**Table 1.**
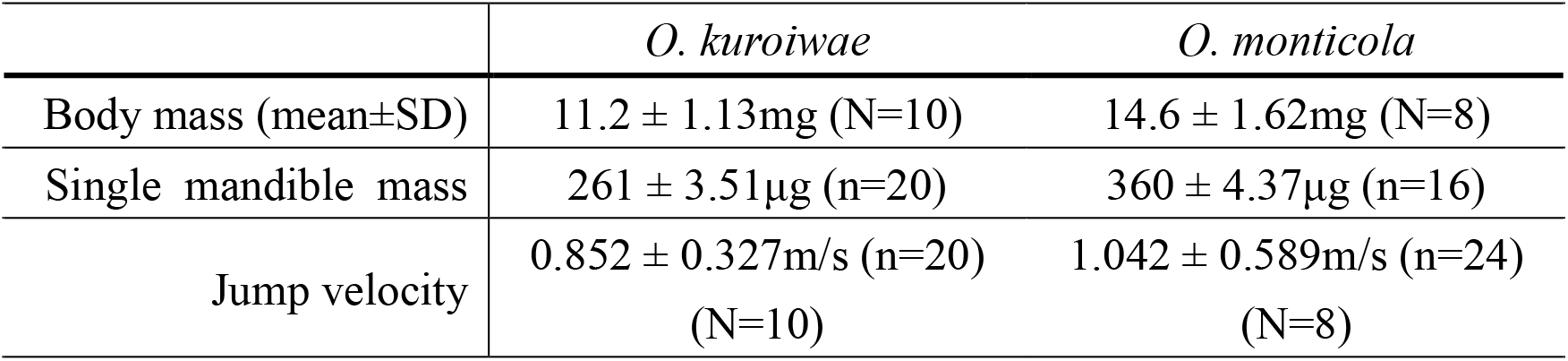
Velocity of mandible powered jump of the trap-jaw ants.

|  | *O. kuroiwae* | *O. monticola* |
| --- | --- | --- |
| Body mass (mean±SD) | 11.2 ± 1.13mg (N=10) | 14.6 ± 1.62mg (N=8) |
| Single mandible mass | 261 ± 3.51μg (n=20) | 360 ± 4.37μg (n=16) |
| Jump velocity | 0.852 ± 0.327m/s (n=20)<br>(N=10) | 1.042 ± 0.589m/s (n=24)<br>(N=8) |

### Mandible powered bouncer defense

Mandible powered jumps (“retrosalient”; Wheeler 1922) were observed during predatorprey encounters (Suppl. Movie S1). To initiate the mandible powered jump, we used the tactile stimulation of the mandible. Since most of the ants responded with retrosalient escape behavior to the unexpected tactile stimulus (Aonuma, 2020), we analyzed this bouncer defense. Ballistic propulsion originated from the ultrafast movement of the mandible (Fig. 1A, B, Suppl. Movie S1). The maximum angular velocity of the mandible was (2.3 ± 0.7) ×10^4^ rad/sec (N=5, n=46, mean ± SD) in *O. kuroiwae* and (1.4 ± 0.4) ×10^4^ rad/sec (N=4, n=38, mean ± SD) in *O. monticola*. Although *O. kuroiwae* closed the mandible significantly faster than *O. monticola*, there was no significant difference in the velocity of the mandible powered jump between *O. kuroiwae* and *O. monticola* (Fig. 1C). The maximum velocity of the mandible powered jump in *O. kuroiwae* was 0.852 ± 0.327 m/sec (N=10, n=20, mean ± SD) and that in *O. monticola* was 1.042 ± 0.589 m/sec (N=8, n=24).

**Figure 1:**
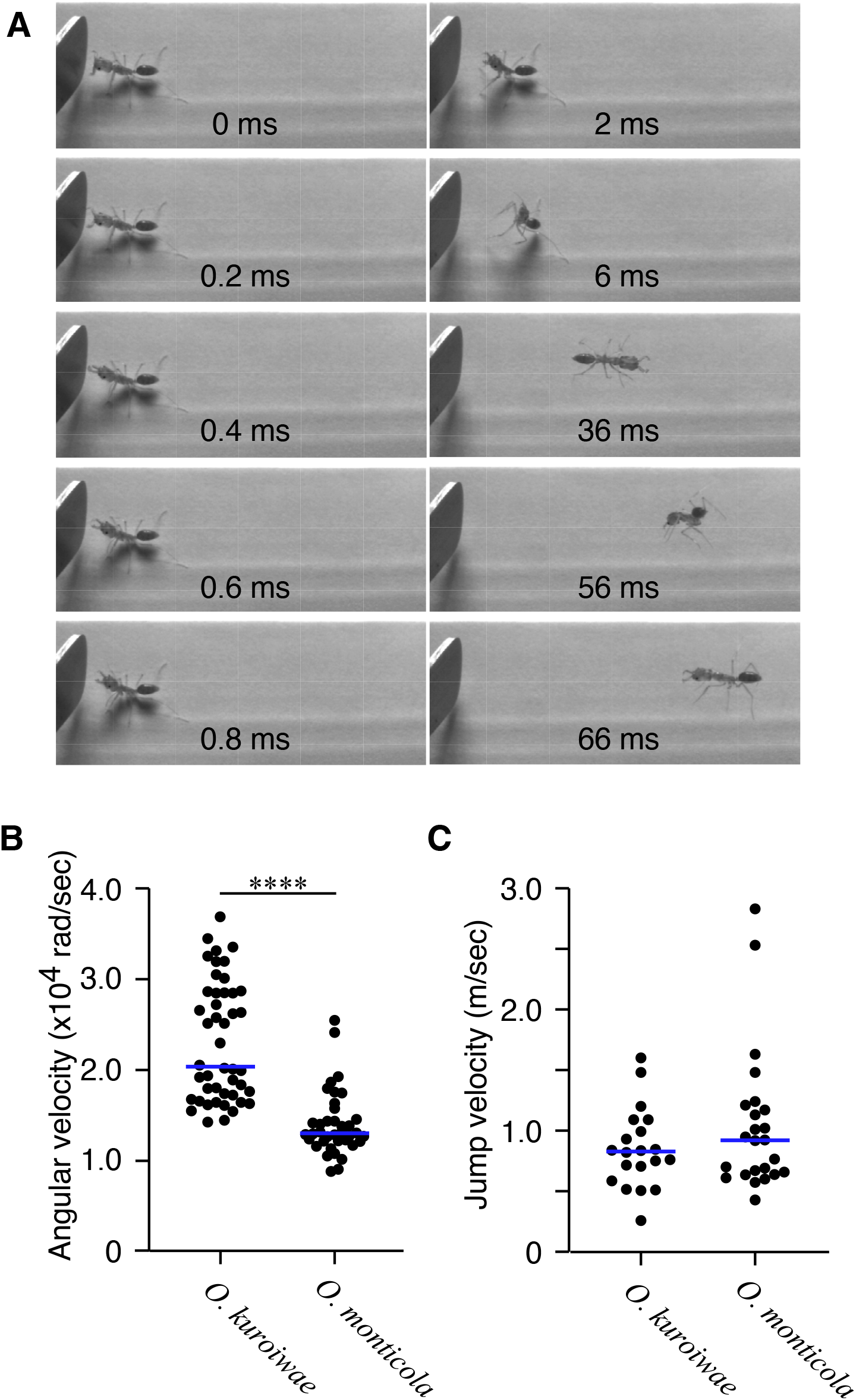
(A) Mandible powered jump of the trap-jaw ant *O. kuroiwae*. The trap-jaw ant shows a defensive posture towards a potential threat. The ant opened the mandible widely before closing them to produce an extremely fast strike to an obstacle (handle of the forceps). The reaction force of the strike made the ant jump. It took within 0.4 msec to close the mandible from the fully opened position to the closed position. The behavior was recorded using a high-speed camera and the recording frame rate was 10,000 fps. (B) Angular velocity of the mandibles during ultra-fast movements of the 2 species *Odontomachus kuroiwae* (n=46) and *Odontomachus monticola* (n=38). The angular velocity of the mandible in *O. kuroiwae* was significantly faster than that in *O. monticola* (p< 0.0001, Mann-Whitney test). Each dot plotted indicates the value of each trial. The bar indicates the median. (C) The maximum jump velocity of the ants. The jump speed of both species was not significantly different (Mann-Whitney test). Each dot plotted indicates the mass of the single side of the mandible. The bar indicates the median.

### Latching the mandible

To power the mandible at an ultra-high speed, trap-jaw ants utilize the latching mechanism of the mandibular joint. To reveal the anatomical structure of the latch, we performed X-ray micro-imaging which allowed us to record the arrangement of mandibular muscles and the three-dimensional anatomical structure of the apodeme, tentorial arm, tentorium, and exoskeleton of the ant (Fig. 2, Suppl. Movie S2). The movement of the mandible is expressed by the sequential movements: open, latch, load, and strike. During the opening phase, contraction of the abductor muscles opened the mandible fully, and then the mandible was kept open until the adductor muscles contracted to close it. The adductor muscles contributed to closing the mandibles both at slow speed and at ultra-high speed. The loading phase, driven by a strong contraction of the adductor muscles, is necessary to power the mandible at an ultra-high speed.

**Figure 2:**
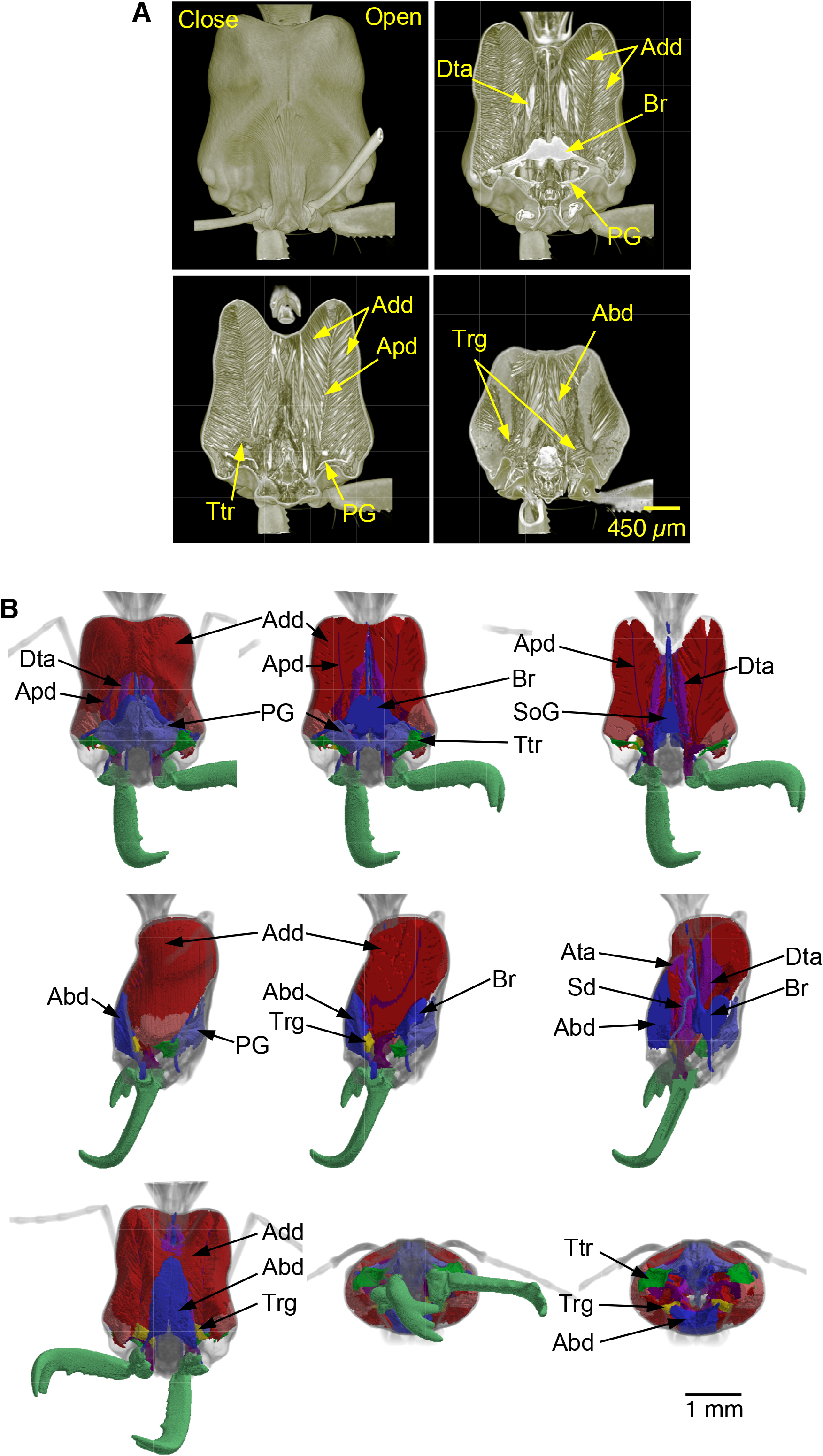
Reconstruction of the X-ray micro-CT imaging of the ant (Sup. 2). (A) Dorsal view of the head that was reconstructed at different focal planes using VGstudio Max Ver 2.2 software. (B) Segmentation images of the adductor, abductor and trigger muscles using amira 3D Ver.2021.1 software. Abd: abductor muscles, Add: adductor muscles, Adt: anterior tentorial arm, Apd: apodeme, Br: brain, Dta: dorsal tentorial arm, PG: pharyngeal gland, Sd salivary duct, SoG: suboesophageal ganglion, Trg: trigger muscle, Ttr: tentorium.

To observe the adductor muscles during the open phase, we CT-scanned the ant with the mandible on one side opened and the other side closed. When the mandible was fully opened, the adductor muscles were extended 157.6± 6.6% (mean ± SD, N=5) compared to the closed side (Table 2, Fig. 2, Suppl. Movie S2). The extension of the adductor muscles was indirectly caused by the contraction of the abductor muscles. The contraction of the abductor muscles rotates the mandible to open, which pulls the apodeme to extend the adductor muscles (Fig. 2A, B). The angle between the muscle fibers and the apodeme on the open side was narrower than on the closed side. The angles between the adductor muscle fibers and apodeme progressively almost narrowed linearly with opening of the mandible (close: 1.07± 0.07 rad (mean ± SD, n=20), half-open: 0.87± 0.04 (n=14), fully open: 0.67± 0.05) (n=16) (Fig. 3A). On the other hand, on the closed side the abductor muscles were more contracted than the open side abductor muscles (Table 2). The micro-CT scanning allowed us to observe the trigger muscles (Fig. 2, 3B). The trigger muscle is thought to contribute to releasing the latched mandible to power at an ultra-high speed (Gronenberg, 1995). The contraction of the abductor muscle also indirectly extended the trigger muscles 135.5± 4.4 % (Table 2). The angle of the trigger muscles to the midline was changed when the mandible was fully opened (Fig. 3C). The angle of the trigger muscles changed with the opening of the mandible (close: 0.96± 0.13, half-open: 1.36± 0.17, fully open: 1.94± 0.16 rad). The change in the angle from a half-open position to a fully opened position was more than that from a closed position to a half-open position. The apodeme attaching to the adductor muscles was pulled with slightly rotating during the opening phase that is caused by the contraction of the abductor muscles (close: 0.37± 0.05, half-open: 0.26± 0.04, fully open: 0.18± 0.03 rad). (Fig. 3D).

**Table 2.** Stretch rate of mandibular muscles (percentage compared to length at rest).

|  | Half-open (N=4) | Entirely open (N=5) |
| --- | --- | --- |
| Adductor muscles | 129.1 ± 4.0 % | 157.6 ± 6.6% |
| Trigger muscles | 128.8± 4.6 % | 135.5± 4.4 % |
|  | Half-close (N=4) | Entirely close (N=5) |
| Abductor muscles | 116.7 ± 4.2 % | 135.4 ± 9.4% |

**Figure 3:**
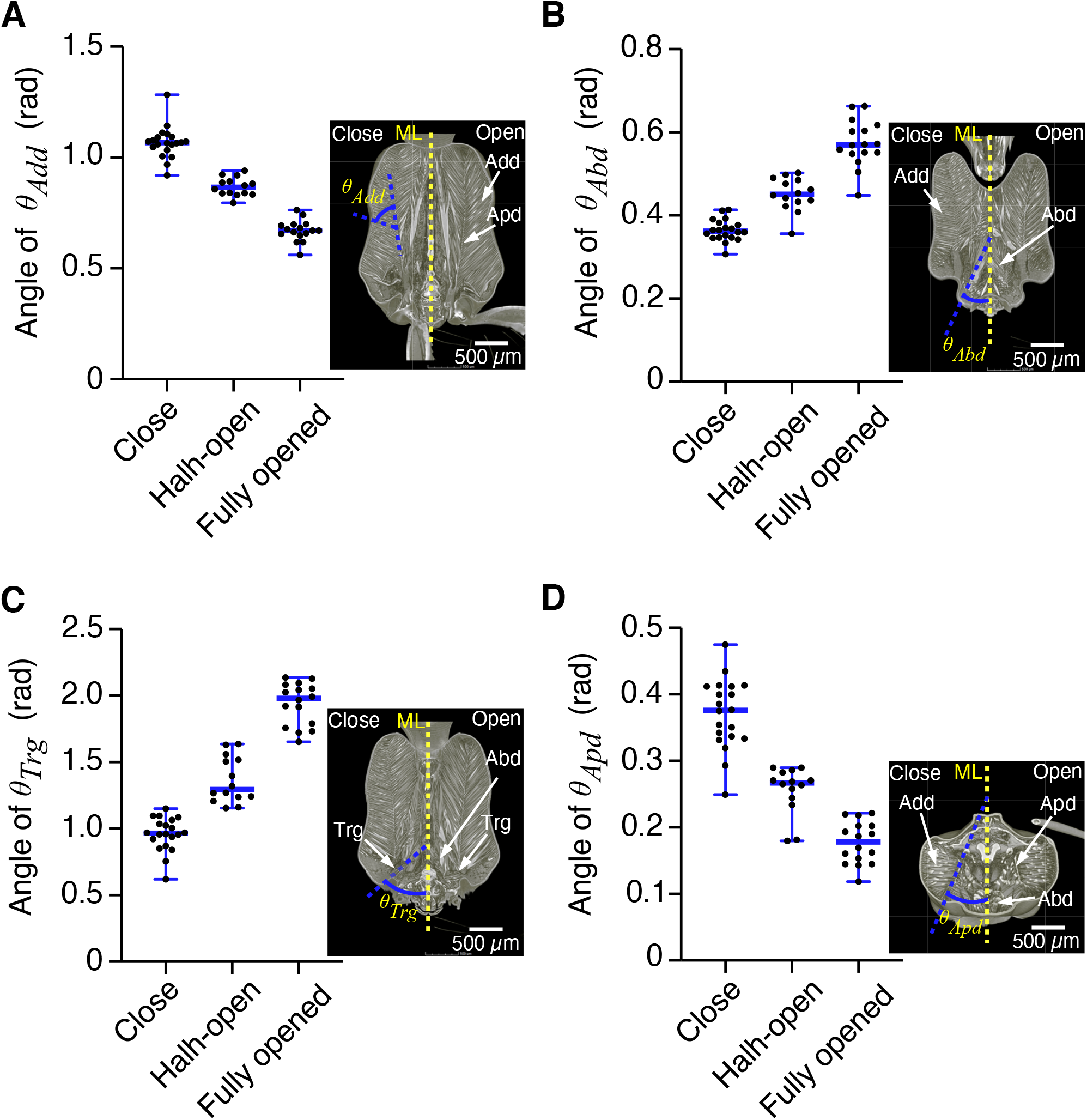
Change in the angles of mandibular muscles and apodeme. (A) Change in the angle of the adductor muscles to apodeme (*θ_Add_*) when the mandible fully opened. (B) Change in the angle of the abductor muscles to midline (*θ_Abd_*) when the mandible fully opened. (C) Change in the angle of the trigger muscles to midline (*θ_Trg_*) when the mandible fully opened. (D). Change in the angle of the apodeme to midline (*θ_Apd_*) when the mandible fully opened. The bars indicate minimum, median and maximum. Each dot plotted indicates the angle of each sample. ML: midline.

The rotation of the mandibular joint also moved the dorsal tentorial arm and tentorium (Fig. 4). The *in vivo* X-ray live imaging showed that the dorsal tentorial arm slightly deformed during the opening phase (Fig. 4B, Suppl. Movie S3). The shape of the dorsal tentorial arm was maintained until the contraction of the adductor muscle occurred to close the mandible. During the loading phase, strong contraction of the adductor muscles deformed the exoskeleton of the head capsule, and the shape of tentorial arm was almost returned to its original shape (Fig. 4C, Suppl. Movie S3). The strike phase is caused by the release of the mandible latch. After striking, the dorsal territorial arm changed slightly to its original shape (Fig. 4D). The head capsule deformation induced by the adductor muscle contraction also returned to the original shape. Live imaging also showed that the anterior tentorial arm was deformed during the opening and loading phase, and then the deformed shapen returned to the original shape after striking (Suppl. Movie S3).

**Figure 4:**
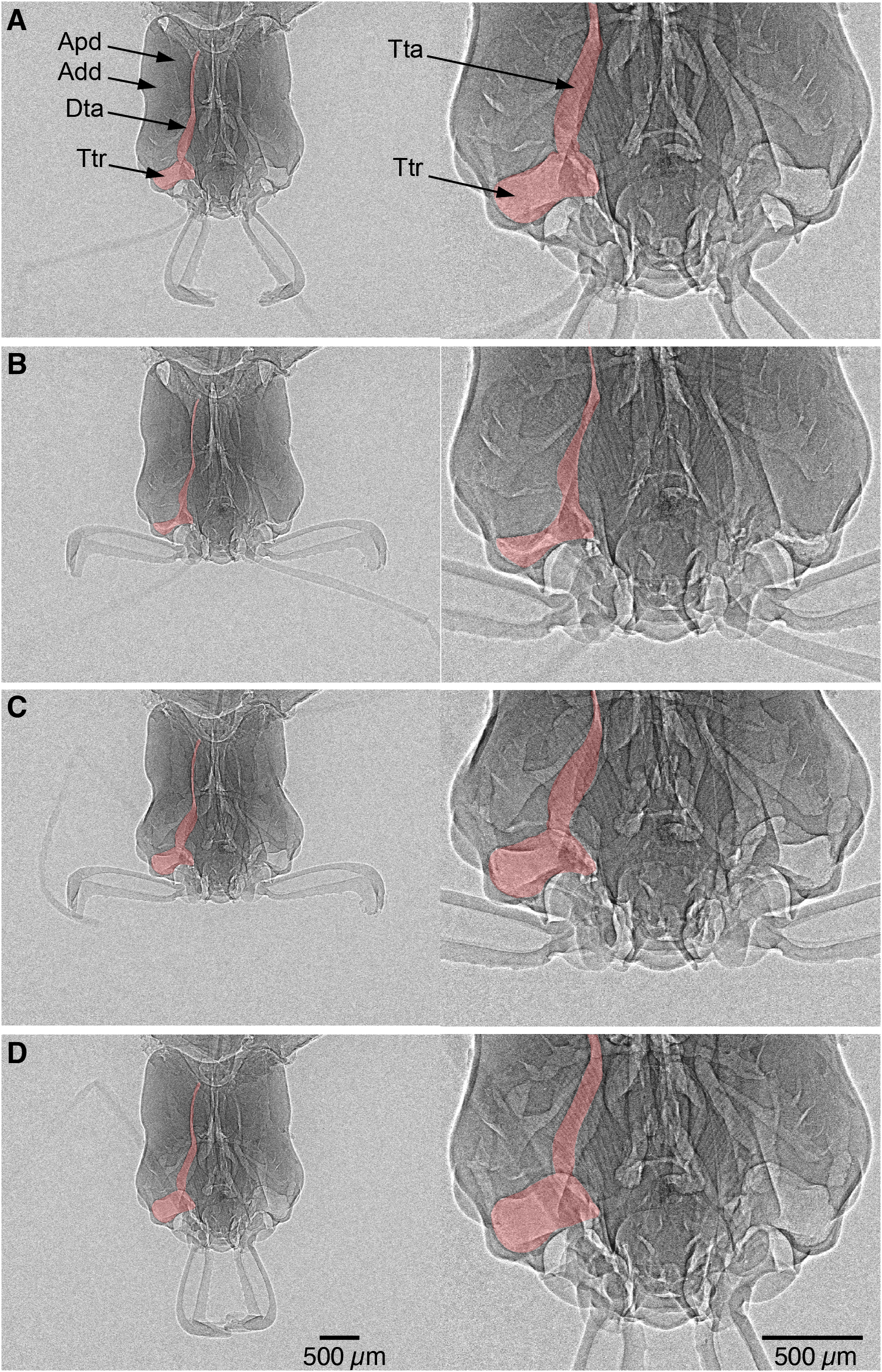
X-ray imaging of the head of a live ant (Suppl. Movie S3). (A) Relaxed position of the mandible. (B) At the end of the open phase, the mandible is fully opened and latched. The dorsal tentorial arm that connects to the adductor muscle was slightly rotated. (C) Loading phase. The exoskeleton of the head capsule deformed by the strong contraction of the adductor muscles. The rotated tentorial arm almost returned to the original position. (D) Strike phase. Just after the strike, the deformed head capsule returned to its original shape. The tentorial arm showed little movement.

The *in vivo* X-ray live imaging demonstrated that the structure of the mandibular joint was a ball-socket like joint (Fig. 5, Suppl. Movie S3). The bulge at the jaw acts as a ball to rotate into the socket when the mandible opens. To identify the anatomical structure of the latch at the mandibular joint, X-ray micro-CT images were then reconstructed (Fig. 6, 8). The micro-CT scanning revealed that there was a U-shaped rail-like structure on the ventral surface of the lip at the socket part of the joint (Fig. 6, Suppl. Movie S2). At the ventral side of the ball, there was a protrusion that did not slide into the socket when the mandible opened but slid on the rail (Fig. 6C, D).The shape of the groove at the ball is in good agreement with the shape to the rail (Fig. 6E, D). These parts were rigidly engaged during the loading phase (Suppl. Movie S4). To examine how the protrusion slides on the rail, the mandibular joint was reconstructed using a 3D printer (Fig. 7, Suppl. Movie S5). The 3D model well reproduces the anatomical structure. The 3D model demonstrated that there is a projection just before the terminal of the rail (Fig. 7B). The protrusion slid on the rail of the socket (Fig. 6E) and the tip of it could get over the projection to terminate at the groove-like structure (Fig. 6, and Fig. 7).

**Figure 5:**
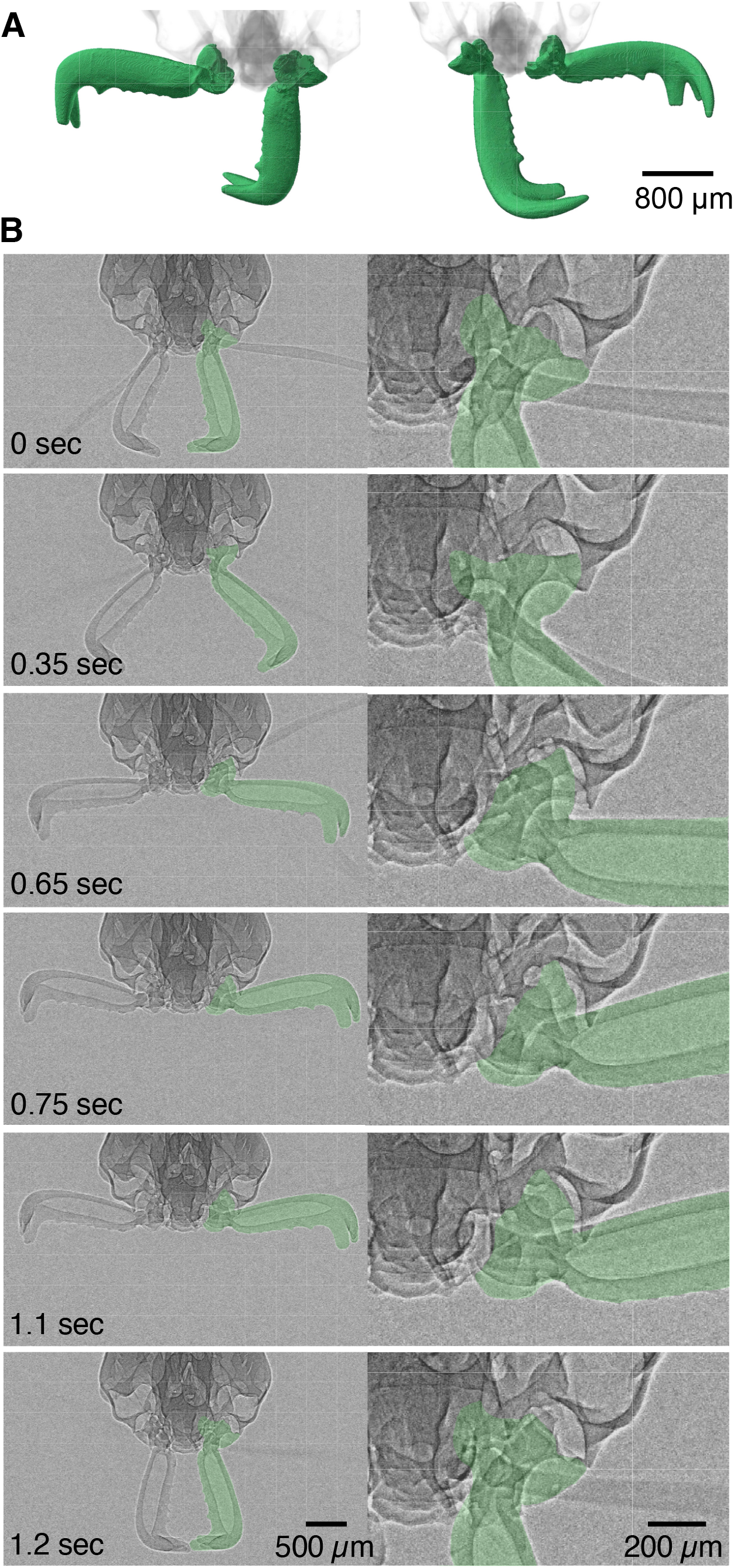
X-ray live imaging of the head of a live ant (Suppl. Movie S3). (A) Reconstruction of the mandible of the ant. The left side is a dorsal view, and the right side is a ventral view of the mandible. (B) Snapshots of the X-ray live imaging during the opening phase (0 - 0.75 sec), loading phase (1.1 sec), and strike phase (1.2 sec).

**Figure 6:**
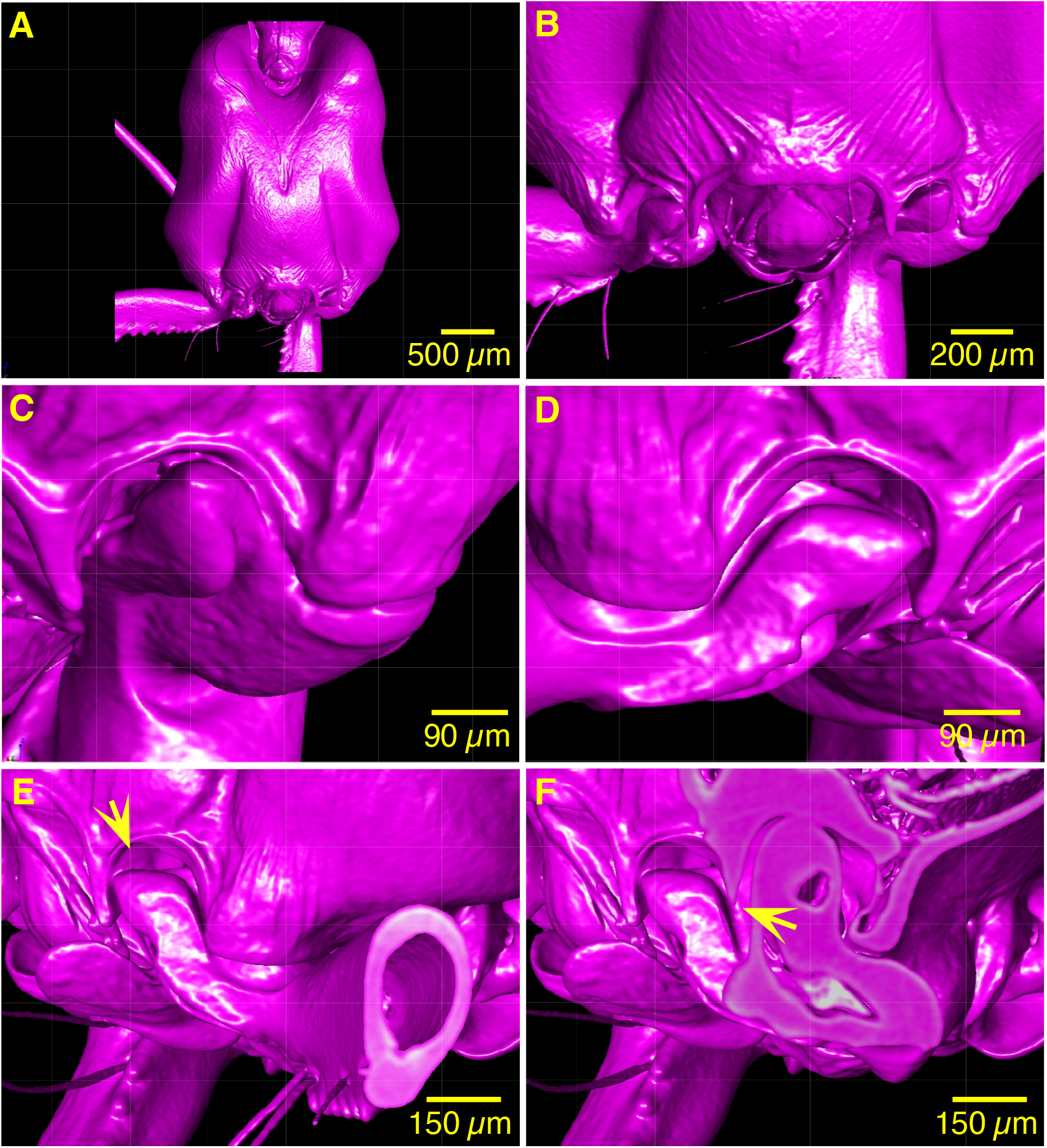
Reconstruction of the X-ray-CT images (Suppl. Movie S2). (A) Ventral view of the head. (B) Ventral view of the mandibular joints. (C) Ventral view of the mandibular joint when the mandible is closed. The protrusion at the ball part of the mandible is outside of the rail-like structure of the ridge of the socket part of the mandibular joint. (D) Ventral view of the mandibular joint when the mandible is fully open. The protrusion at the ball part is on the rail. (E) Lateral view of the mandibular joint when fully open. The arrow indicates the projection on the surface of the rail. (F) Sagittal view of the rail indicated by arrow.

**Figure 7:**
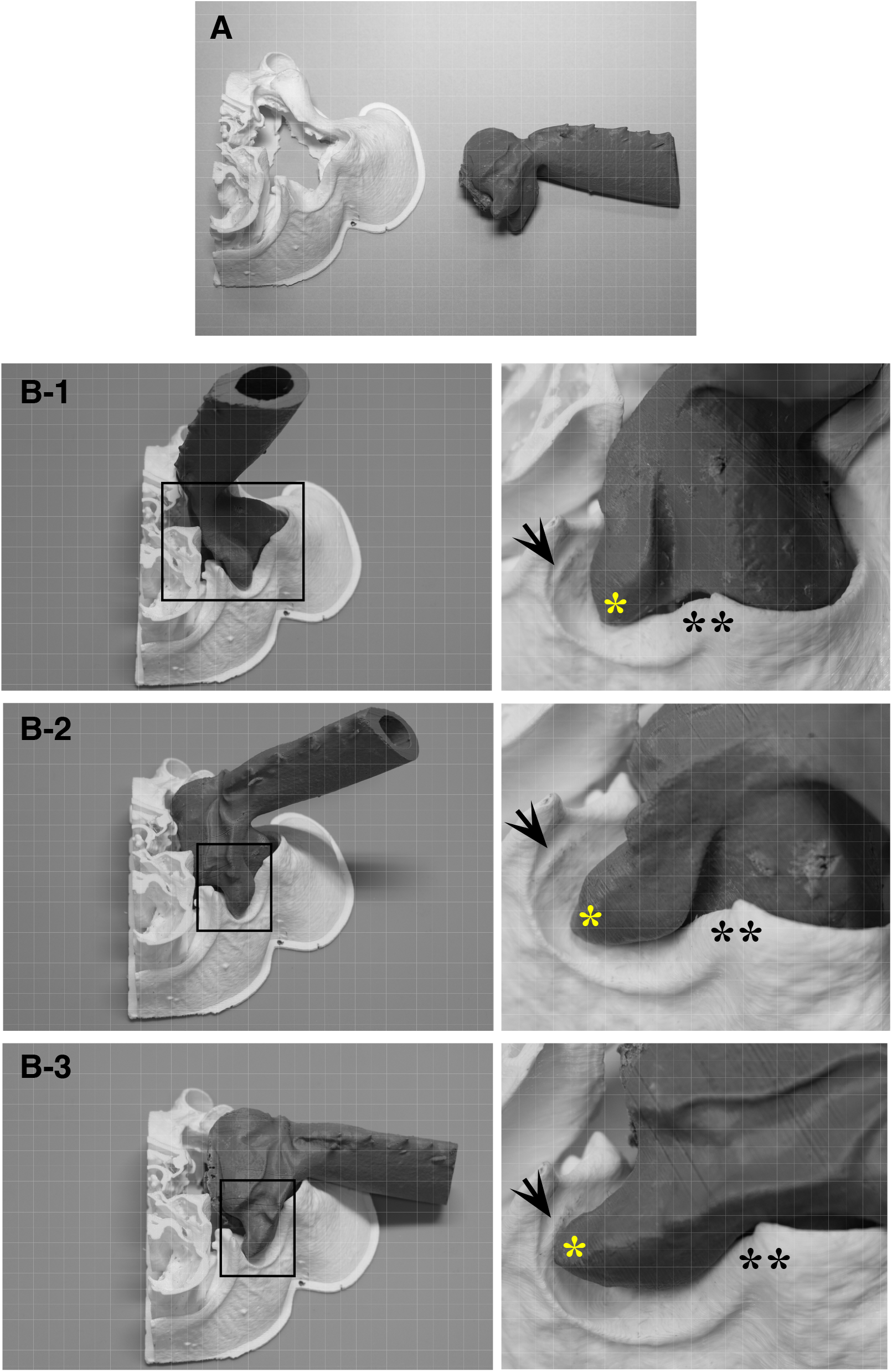
3D model of the mandibular joint (Suppl. Movie S5). (A) Socket and ball of the mandibular joint were separately printed out using a 3D printer. (B) The ball of the joint was placed in the socket and moved. From B-1 to B-3 are snapshots of the different angles of the joint. Right-side images are the rail system of the mandible that is framed in the area indicated in the left-side images. The single asterisk indicates the protrusion at the ball part of the mandible. Double asterisks indicate the rail-like structure of the ridge of the socket part of the mandibular joint. Arrow indicates another projection where the protrusion of the ball is terminated.

**Figure 8:**
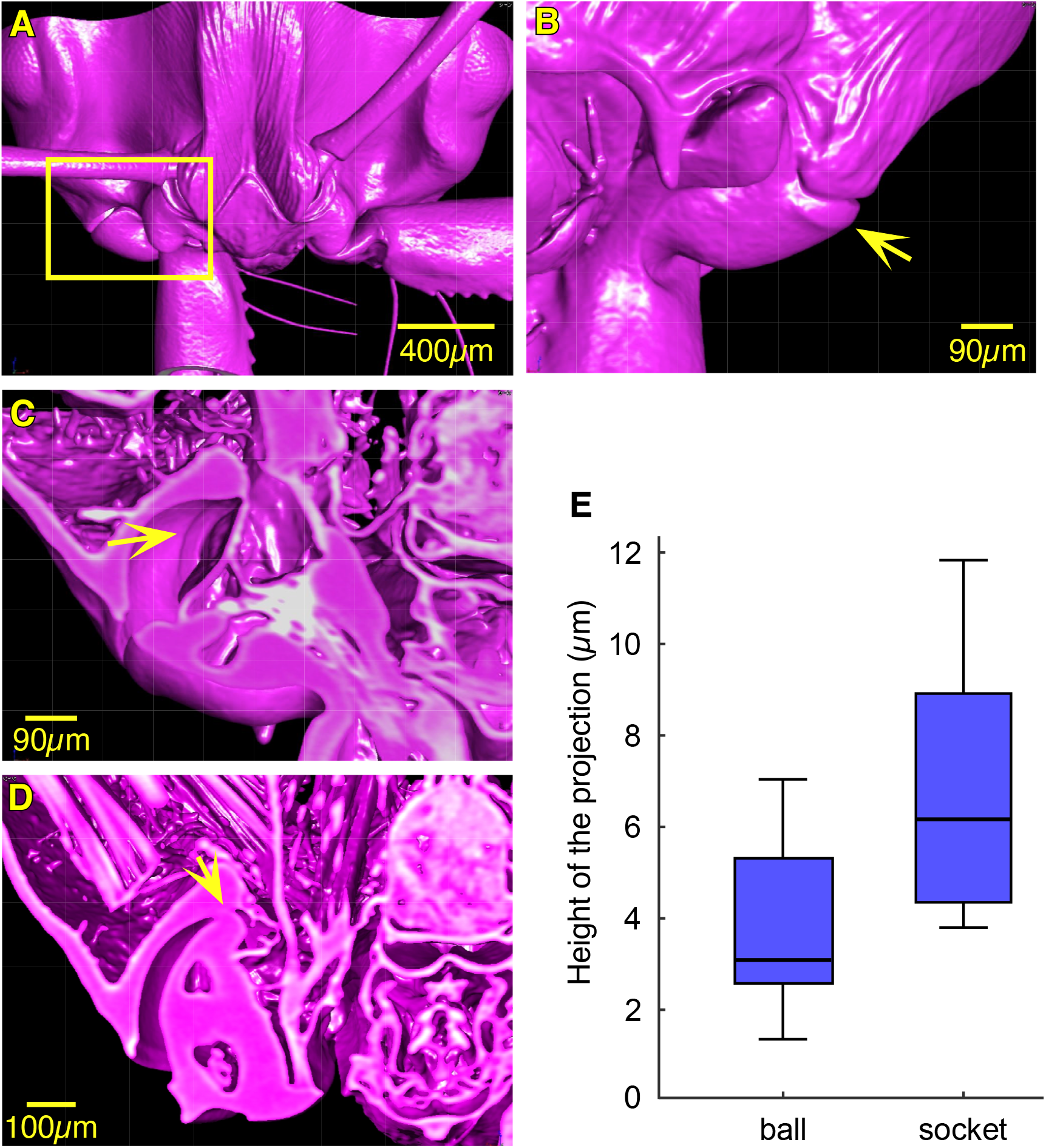
(A) Reconstructed images of the mandibular joint (Suppl. Movie S2). (B) Ventral view of the framed area indicated in A. The arrow shows the projection of the ball part of the mandibular joint. (C) Inside the socket part of the mandibular joint when the mandible is close. The arrow indicates the projection on the surface of the exoskeleton. (D) Inside of the socket part of the mandibular joint when the mandible is fully open. The arrow shows the rigidly engaged part of the projection of the ball and the groove of the socket. (E)The height of the projection on the surface of the mandibular joint. Box-and-whisker graphs indicate minimum, median, maximum, and 25th and 75th percentiles.

Micro-CT scanning revealed that the ball of the mandible had a fine projection on the surface of the ball lip (Fig. 8B, arrow). The height of the projection was 3.8 ± 1.8 μm (N=7, n=12, mean ± SD) and this part slides into the socket of the joint when the mandible is entirely opened (Fig. 5). Micro-CT scanning also revealed that there was another projection on the surface of the inner section of the socket. The height of that projection on the surface of the cuticle at the socket was 6.8 ± 2.7 μm (N=7, n=12, mean ± SD). These anatomical structures of the mandible ball joint suggest that the ball that has a projection slides into the socket and over the projection at the socket followed by sliding motion into the groove (Fig. 8D). If this is the case, the angular velocity of the mandible during the open phase must change when the ball slides over the projection and into the groove.

### Latching kinematics

To confirm quantitatively the functional aspect of the projection on the surface of the ball corresponding to the groove at the end of the socket, we examined the change in the angular velocity of the mandible during the opening phase (Fig. 9). The movements of the tip of the mandible were tracked and analyzed (N=10, n=42). The latching kinematics demonstrated that the angular velocity of the mandible increased just before the fully opened position. The peak in the angular velocity occurred approximately 0.056 (π/18) rad before the final position of the mandible. This movement indicates that the mandible was sliding into the final stable position. The blue line in the Figure 9A indicates the estimated angular velocity as a function of angular position of the mandible based on the statistical model (see Materials and Methods). We constructed two statistical models to examine possible kinematic differences between the two ant species, *O. kuroiwae* and *O. monticola*, however very similar kinematic changes were obtained for both species. Variations among the trials and individuals were larger than those of species. Since the anatomical structure of the ball joint of the two species were also quite similar, we show the result estimated from the statistical models of both species combined.

**Figure 9:**
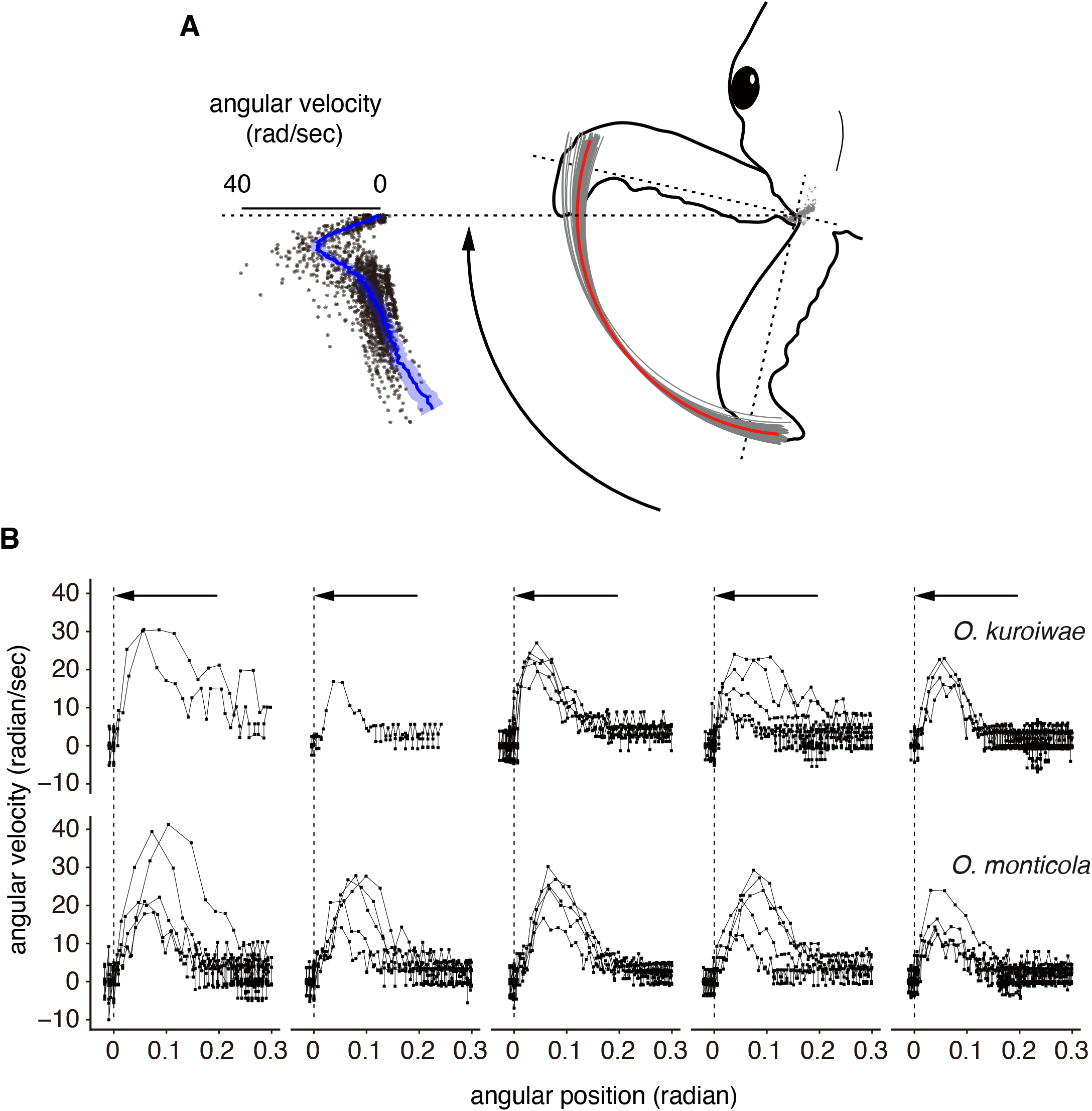
Latching kinematics indicate that the opening velocity of the mandible increases just before reaching the latched position. (A) The ants open the mandible, and the velocity of the movement shows a peak at approximately π/18 rad before the final position of the mandible, revealing that the mandible is levered into the final stable position. The blue line marks the estimated angular velocity as a function of the angular position of the mandible based on the statistical models (see Materials and Methods). (B) Change of the opening velocity for each individual and species. It largely varied within and across individuals, but no difference between species.

## Discussion

Trap-jaw ants strike using their mandibles with an ultra-high speed when hunting (De la Mora et al., 2008) and during mandible-powered jumping (Larabee and Suarez, 2015). The genus *Odontomachus* provided a good model system to investigate the mechanisms underlying these ultra-fast movement of the appendage (Gronenberg et al., 1993; Larabee and Suarez, 2015; Sutton et al., 2022; Wang et al., 2020). The maximum angular velocities of the mandible have been reported in some *Odontomachus* species (Spagna et al., 2008; Spagna et al., 2009; Larabee et al., 2017). Here, we gained information about other two species, *O. kuroiwae* and *O. monticola*. Although the maximum angular velocity of *O. kuroiwae* was higher than that of *O. monticola*, the velocity of the bouncer defense was similar. Since the body mass of *O. kuroiwae* was smaller than that of *O. monticola*, *O. kuroiwae* may be more influenced by the viscosity of the air during the jump than *O. monticola*. To understand the difference in the velocity between the two species, it was necessary to measure the strike force of the mandibles and kinematic analysis of the mandible powered jump is necessary as a next step.

Prior to the mandible strike, the trap-jaw ants load elastic energy and amplify the power to produce ultra-fast movements (Patek et al., 2006; Sutton et al., 2022). During the loading, strong contraction of the mandible adductor muscles pulls the adductor apodemes and deforms the exoskeleton of the head and tentorial arms. The elastic deformation of the apodeme and skeleton function as a spring to store the energy to produce the power for striking.

Spring and latch systems, including artificial systems, are diverse and operate under their own mass-specific power limits (Ilton et al., 2018). In the locust, the semi-lunar process (SLP) at the femur-tibia joint is bent to store energy for use during kicking and jumping (Cofer et al., 2010). To release amplified power to produce extremely fast movements, catapult mechanisms are thought to be the key in invertebrates (Wan et al., 2016, Bolmin et al., 2019, Ritzmann, 1973; Ritzmann, 1974; Kaji et al., 2018, Patek et al., 2004; Patek et al., 2007; Zack et al., 2009) The catapult mechanism is also thought to contribute to producing quick movement in vertebrates, such as in the frog jump (Astley and Roberts, 2012; Astley and Roberts, 2014). To utilize catapult mechanisms, arthropods have evolved diverse latch mechanisms to lock the powered body part in place while building up elastic energy to be released as kinetic energy during their strike (Ilton et al., 2018).

Releasing a latch allows a joint to move extremely quickly via recoil. The mechanisms underlying ultra-fast movements of the trap-jaw ants have been investigated and the latch mechanism to lock the joint has been demonstrated previously (Gronenberg, 1995), however, the detailed anatomical structure of the latch remained unclear. The X-ray live imaging and 3D model allowed us to observe the structure and the movement of the mandibular joint. The joint of the mandible forms a ball joint that could make the mandible movement stable. The micro-CT scanning revealed a U-shaped rail structure on the ventral surface of the lip of the socket part of the joint that could also stabilize mandible movements. The rail and protrusion consist of the hard cuticle and these anatomical structures function to rotate the mandible stably open. The anatomical structure consists of a protrusion tip and a groove after the projection that could function to lock the mandible rigidly. The hard cuticle of these structures could be a support for the ball, preventing it from slipping into the socket during the loading phase and deformation of the elastic exoskeletal structure of the head. These anatomical structures therefore not only function as a part of the latch system but also function to support the stability of the mandible during a strike.

X-ray micro-CT scanning revealed another potential anatomical candidate that forms part of the latch system in the mandibular joint. The ball of the mandible has a projection at the lip of the lateral area and when the mandible opens, this area slides into the socket of the mandibular joint. The socket also has a projection of a similar size. These anatomical structures can function as a detent ridge, i.e., the ball slides into the socket and the projection of the ball gets over the projection of the socket followed by sliding into the groove. The kinematic analysis demonstrated that the angular velocity of the mandible becomes faster just before terminating the movement at the open phase. This detent ridge would also function as a latch mechanism. There are, therefore, at least two latch mechanisms on the mandibular joint; one is on the rail and the other is inside the socket.

During the loading phase, strong contraction of the adductor muscles induces elastic energy storage in the apodeme and skeletal systems, which gradually increases frustration in the multiple spring and latch mechanisms. This critical point, or state, is minimally stable, which makes it sensitive to release by the latch trigger system. The trigger muscles are known to be activated just before the strike (Gronenberg, 1995). Contraction of the trigger muscle applies lateral force to the apodeme, which initiates the mandible strike. Micro-CT scanning demonstrated that the trigger muscle changed angle when abductor muscles contract to open the mandible. The angle to the midline changes from about 1 rad (mandible closed) to 2 rad (fully opened) indicating that the trigger muscle can apply lateral force to the apodeme. Furthermore, the movement of the 3D model demonstrated that the trajectory of the opening mandible slightly twisted just before latching. Since the latching of the mandible is minimally stable a small force applied by the contraction of the trigger muscle would be sufficient to release the latches.

This study demonstrates that the mandibular joint of the trap-jaw ant genus *Odontomachus* has multiple latches forming a system consisting of the detent ridge structures on the ball and the groove inside of the socket, and a rail at the lip of the socket and protrusion at the ball. The morphological structure of arthropod animals is diverse in their appendages and similarly in their joints. Unlike artificial ball joints, the threedimensional structure of the joints in arthropod animals are complex. Indeed, the stricture of the ball part of the mandibular joint is complex. The trap-jaw ants have evolved a ball joint that has a rail structure to move the mandible stably when striking. Multi-latch system would be effective to catch the joint that has a protrusion structure when opening the mandible. Multi-latch system could lock rigidly the mandible at different part of the joint during the loading phase. The height of the projections of the latch system are 3-5 μm, which make each latch minimally stable. Small displacements caused by the contraction of the trigger muscles could change the balance of the minimally stable joint to release the latches. To further gain an understanding of the mechanisms to power the mandible at an ultra-high speed, it will be necessary to investigate how each mandibular muscle contributes to maintain the stability of the multi latch system and break the balance to release the latch. *In vivo* X-ray diffraction live imaging will allow us to investigate this in detail in the future.

## Acknowledgements

A part of this work was supported by JST CREST Grant Number JPMJCR14D5, and by Japan Society for the Promotion of Science KAKENHI [Grant-in-Aid for Scientific Research (S), grant JP17H06150, Grant-in-Aid for Scientific Research (A), grant number 22H00216, and Grant-in-Aid for Challenging Exploratory Research, grant number 22K19795], Japan. Synchrotron experiments were supported by JASRI (2018A1240 and 2018A1244). We thank Drs. K. Uesugi and H. Iwamoto in Spring-8 for their comments and technical support. We are also grateful to Prof. Philip L. Newland at the University of Southampton in the U.K. and Ms. Valeria Zeni at the University of Pissa in Italy for their critical reading and comments on the manuscript.

## Supplementary

**Suppl. Movie S1:** Highspeed movie of mandible powered jump captured at 10,000 fps.

**Suppl. Movie S2:** Movie of the X-ray micro-CT images of the trap-jaw ant.

**Suppl. Movie S3:** X-ray live imaging during the ultra-fast movement of the mandible of the trap-jaw ant.

**Suppl. Movie S4:** 3D model of the mandibular joint. Digital data of the micro-CT images were converted into STL files and then printed out using a 3D printer. The movie was made using a series of snapshot of the 3D model with different positions of the mandible.

**Suppl. Movie S5:** Movement of the mandibular joint was recorded from the ventral side. During the loading phase, the rail and the groove of the ball were rigidly engaged.

## Notes

### Competing Interest Statement

The authors have declared no competing interest.

### Summary of Updates

Some descriptions and figures were improved.

